# A broadly neutralizing biparatopic Nanobody protects mice from lethal challenge with SARS-CoV-2 variants of concern

**DOI:** 10.1101/2021.08.08.455562

**Authors:** Teresa R. Wagner, Daniel Schnepf, Julius Beer, Karin Klingel, Natalia Ruetalo, Philipp D. Kaiser, Daniel Junker, Martina Sauter, Bjoern Traenkle, Desiree I. Frecot, Matthias Becker, Nicole Schneiderhan-Marra, Annette Ohnemus, Martin Schwemmle, Michael Schindler, Ulrich Rothbauer

## Abstract

The ongoing COVID-19 pandemic and the frequent emergence of new SARS-CoV-2 variants of concern (VOCs), requires continued development of fast and effective therapeutics. Recently, we identified high-affinity neutralizing nanobodies (Nb) specific for the receptor-binding domain (RBD) of SARS-CoV-2, which are now being used as biparatopic Nbs (bipNbs) to investigate their potential as future drug candidates. Following detailed *in vitro* characterization, we chose NM1267 as the most promising candidate showing high affinity binding to several recently described SARS-CoV-2 VOCs and strong neutralizing capacity against a patient isolate of B.1.351 (Beta). To assess if bipNb NM1267 confers protection against SARS-CoV-2 infection *in vivo*, human ACE2 transgenic mice were treated by intranasal route before infection with a lethal dose of SARS-CoV-2. NM1267-treated mice showed significantly reduced disease progression, increased survival rates and secreted less infectious virus via their nostrils. Histopathological analyses and *in situ* hybridization further revealed a drastically reduced viral load and inflammatory response in lungs of NM1267-treated mice. These data suggest, that bipNb NM1267 is a broadly active and easily applicable drug candidate against a variety of emerging SARS-CoV-2 VOCs.

## Introduction

The ongoing SARS-CoV-2 pandemic continues to be challenging due to limited access to vaccines in certain countries or vaccine fatigue in others, the lack of effective and easy-to-administer antivirals, and the emergence of variants of concern (VOCs) (Scudellari, 2020). Despite the rapid development of effective vaccines, global immunity or alternatively eradication of SARS-CoV-2 is currently out of reach (Dagan *et al*, 2021; Kwok *et al*, 2020). In addition, vaccination does not confer sterile immunity against SARS-CoV-2 infection and especially in the elderly, immunocompromised individuals, or individuals with severe preexisting conditions; breakthrough infections can still develop into life-threatening disease (Beaudoin-Bussières *et al*, 2020; Kustin *et al*, 2021; Long *et al*, 2020). In particular, some SARS-CoV-2 variants with increased transmissibility and pathogenicity accompanied by a partial immune escape were reported to cause severe disease progression even in vaccinated individuals (Becker *et al*, 2021; Challen *et al*, 2021; Davies *et al*, 2021a; Davies *et al*, 2021b; Jewell, 2021; Madhi *et al*, 2021; Volz *et al*, 2021; Zhou *et al*, 2021). Consequently, there is a continuing and urgent need for effective and easy to administer antivirals against emerging SARS-CoV-2 variants. Neutralizing monoclonal antibodies (Nabs) have been granted emergency use authorization by the U.S. Food and Drug Administration and were shown to efficiently reduce mortality in COVID-19 patients with increased risk for a severe disease progression (Chen *et al*, 2020; Jiang *et al*, 2020; Weinreich *et al*, 2020). Most of these Nabs target the interaction site between receptor-binding domain (RBD) of the SARS-CoV-2 spike protein and angiotensin-converting enzyme (ACE) 2 to prevent viral entry into epithelial cells of the respiratory tract (Brouwer *et al*, 2020; Cao *et al*, 2020; Ju *et al*, 2020). However, viral escape from neutralizing antibodies resulted in several mutations affecting the RBD:ACE2 interface, which impairs binding of established NAbs and thus limits current direct-acting antiviral treatment options (Diamond *et al*, 2021; Wang *et al*, 2021).

In parallel to conventional antibodies, camelid single-domain antibody fragments, better known as nanobodies (Nbs), have been developed to target the RBD of SARS-CoV-2 (Chi *et al*, 2020; Hanke *et al*, 2020; Huo *et al*, 2020; Wagner *et al*, 2021; Wrapp *et al*, 2020). Due to the unique physicochemical properties of Nbs such as their small size, stable folding, and efficient tissue penetration, these molecules are considered to be ideal for therapeutic application. Indeed, some of these Nbs showed strong neutralizing efficacy against SARS-CoV-2, especially when used in the multivalent or multiparatopic format (Koenig *et al*, 2021; Nambulli *et al*, 2021; Schepens *et al*, 2021; Wagner *et al.*, 2021; Xiang *et al*, 2020).

Recently, we reported the identification of several Nbs demonstrating a high neutralizing capacity against SARS-CoV-2 and generated a biparatopic (bip) Nb (NM1267) that binds two distinct sites, one epitope inside and one outside of the RBD:ACE2 interface (Wagner *et al.*, 2021). By the application of NM1267 advanced diagnostic assays were developed, determining the emergence of a neutralizing immune response in convalescent or vaccinated individuals (Becker *et al.*, 2021; Wagner *et al.*, 2021). In this study, we generated in addition to NM1267 two novel bipNbs based on Nb pairs targeting different epitopes within the RBD. Upon analyzing their affinities and stabilities in accelerated aging assays, we identified NM1267 as the most promising candidate. Based on these results, we tested the neutralizing potency of NM1267 for a B.1.351 (Beta) patient isolate in direct comparison to SARS-CoV-2 WT in an *in vitro* virus neutralization test (VNT) and determined its protective efficacy *in vivo*, using transgenic mice expressing human ACE2 (K18-hACE2 mice) (McCray Jr *et al*, 2007; Winkler *et al*, 2020). Consistent with its neutralizing activity *in vitro*, NM1267 efficiently protected mice from lethal challenge with SARS-CoV-2 VOCs and profound lung tissue damage, confirming its suitability as promising drug candidate.

## Results

Following our recently reported approach in which we combined two Nbs targeting different epitopes within the RBD of SARS-CoV-2 to generate the strongly neutralizing biparatopic Nb (bipNb) NM1267 (Wagner *et al.*, 2021), we designed two additional bipNbs. We genetically coupled Nbs NM1230 and NM1228, which both bind within the RBD:ACE2 interface, and NM1228 and NM1226, of which the latter targets an epitope outside the RBD:ACE2 interface, via a flexible Gly-Ser ((G_4_S)_4_) linker, resulting in the bipNbs NM1266 and NM1268, respectively (**Supplementary Table 1**). Similar to NM1267 (Wagner *et al.*, 2021), both novel bipNbs were produced with a high yield and good purity in mammalian cells and showed picomolar affinities to wild-type (WT) RBD as measured by biolayer interferometry (BLI) (**Figure 1A**). To test their potency to block the interaction between RBD, S1, or spike of SARS-CoV-2 to human ACE2, we performed a recently established multiplex ACE2 competition assay (Wagner *et al.*, 2021). The results showed that both bipNbs inhibited binding of ACE2 to all tested antigens in the low picomolar range (**Supplementary Figure 1A, C**). Additionally, we confirmed their neutralizing effect in a VNT using SARS-CoV-2 WT which revealed half maximal inhibitory concentrations (IC_50_s) in the nano- or picomolar range (**Supplementary Figure 1B**). Notably, NM1267 proved to be the most potent bipNb in this assay with an IC_50_ of 380 pM (**Supplementary Figure 1C**). For further selection, we next assessed the biophysical properties of all three bipNbs by measuring thermal unfolding and aggregation with nano differential scanning fluorimetry (nanoDSF) **(Figure 1B**). While the bipNbs NM1267 and NM1268, both containing the Nb NM1226, showed a slight increase in light scattering, indicating a higher aggregation tendency at higher, non-physiological temperatures, NM1266 showed no onset of aggregation up to 90°C. Reanalysis after accelerated aging at 37°C for ten days revealed no considerable differences compared to baseline (**Figure 1B**). From these data, we concluded that all three bipNbs are highly stable. However, as NM1267 showed the highest melting temperature of ∼57°C (**Figure 1B**), indicating improved thermal stability, we decided to proceed with the bipNb NM1267 as our favorite candidate for the intended *in vivo* application.

**Figure 1.**
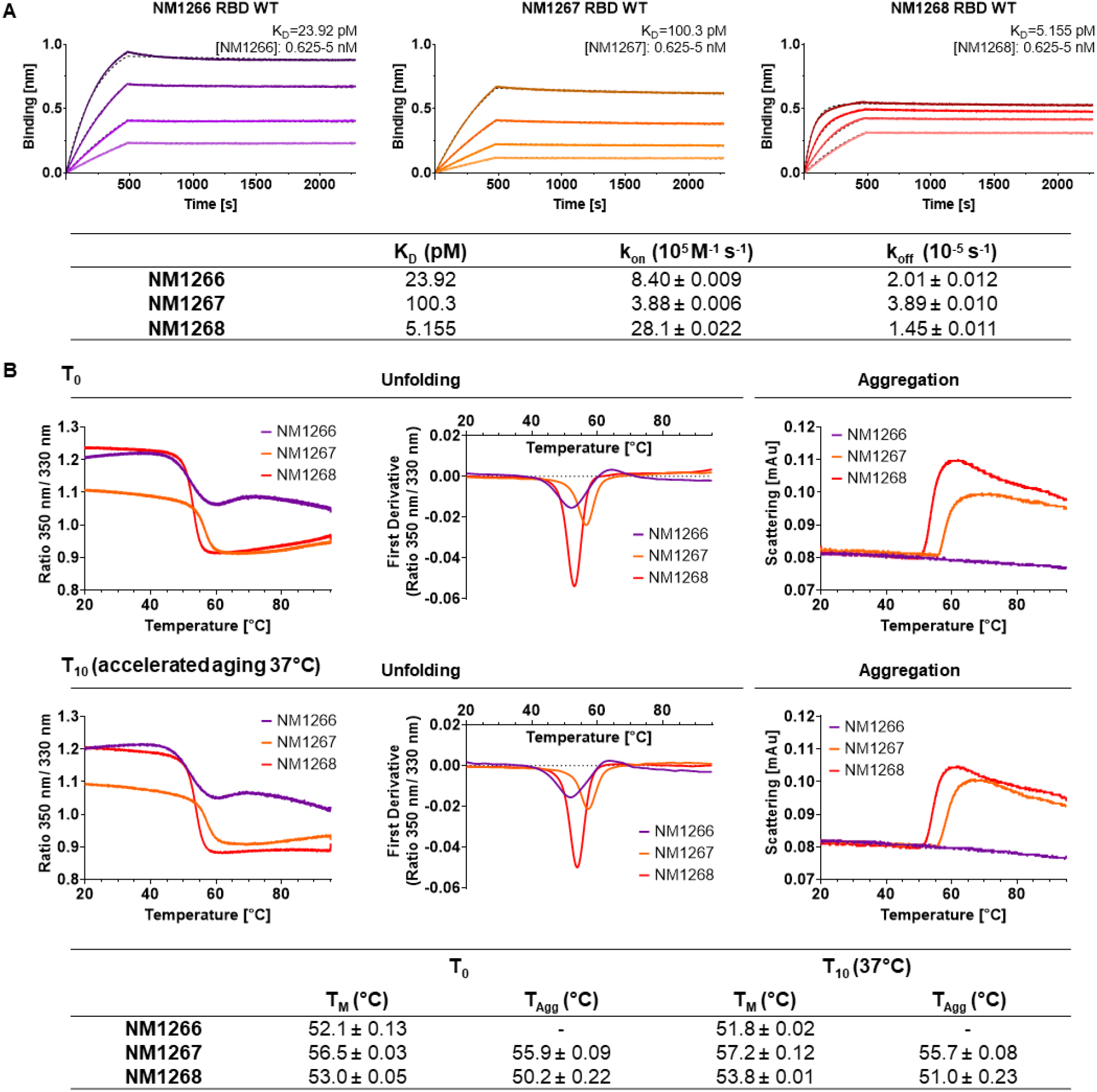
Affinity and stability of different biparatopic Nanobodies. **A** Affinity measurements by biolayer interferometry (BLI) of bipNbs NM1266, NM1267 and NM1268. bipNbs were applied with concentrations ranging from 5–0.625 nM (illustrated with gradually lighter shades) on immobilized wild-type RBD (RBD WT). Global 1:1 fits are illustrated as dashed lines and binding affinity (K_D_), association (k_on_) and dissociation constant (k_off_) determined for the individual bipNbs are summarized. **B** Stability analysis of bipNbs NM1266, NM1267 and NM1268 was performed at time points T_0_ and T_10_ after storage at 37°C for ten days to induce accelerated aging. Protein unfolding was determined by fluorescence emission wavelengths shifts illustrated as fluorescence ratios (350 nm/ 330 nm) and its first derivative. Protein aggregation status was measured by light intensity loss due to scattering. Melting (T_m_) and aggregation (T_Agg_) temperature are summarized as table for both time points.

Previously, we had shown that the individual Nbs linked to form NM1267 (NM1230 and NM1226) both exhibit strong neutralization potency regardless of whether they bind within the RBD:ACE2 interface, as shown for NM1230, or bind a more conserved region outside this interaction site, as shown for NM1226 (Wagner *et al.*, 2021) (**Supplementary Figure 2**). With the precise epitopes known, we considered that NM1267 could also be effective against lately described VOCs, since only mild binding interference were expected due to the acquired mutations (**Supplementary Figure 2**). Therefore, we analyzed binding affinities of NM1267 towards RBDs of emerging SARS-CoV-2 variants using BLI (**Figure 2A-I**). Compared to RBD_wt_, NM1267 showed similar or even better binding to RBDs from B.1.1.7 (Alpha) (**Figure 2A**), B.1.351 (Beta) (**Figure 2B**), P1 (Gamma) (**Figure 2C**), P3 (Theta) (**Figure 2F**) and A.23.1 (**Figure 2H**), while a slight decrease in affinity was observed for RBDs from B.1.617.2 (Delta) (**Figure 2D**), B.1.429 (Epsilon) (**Figure 2E**) and B.1.617.1 (Kappa) (**Figure 2G**). However, the measured affinities in the picomolar range confirmed the high potential of NM1267 to also bind SARS-CoV2 variants with mutations at the RBD:ACE2 interface (**Figure 2A-I, Supplementary Figure 2**).

**Figure 2.**
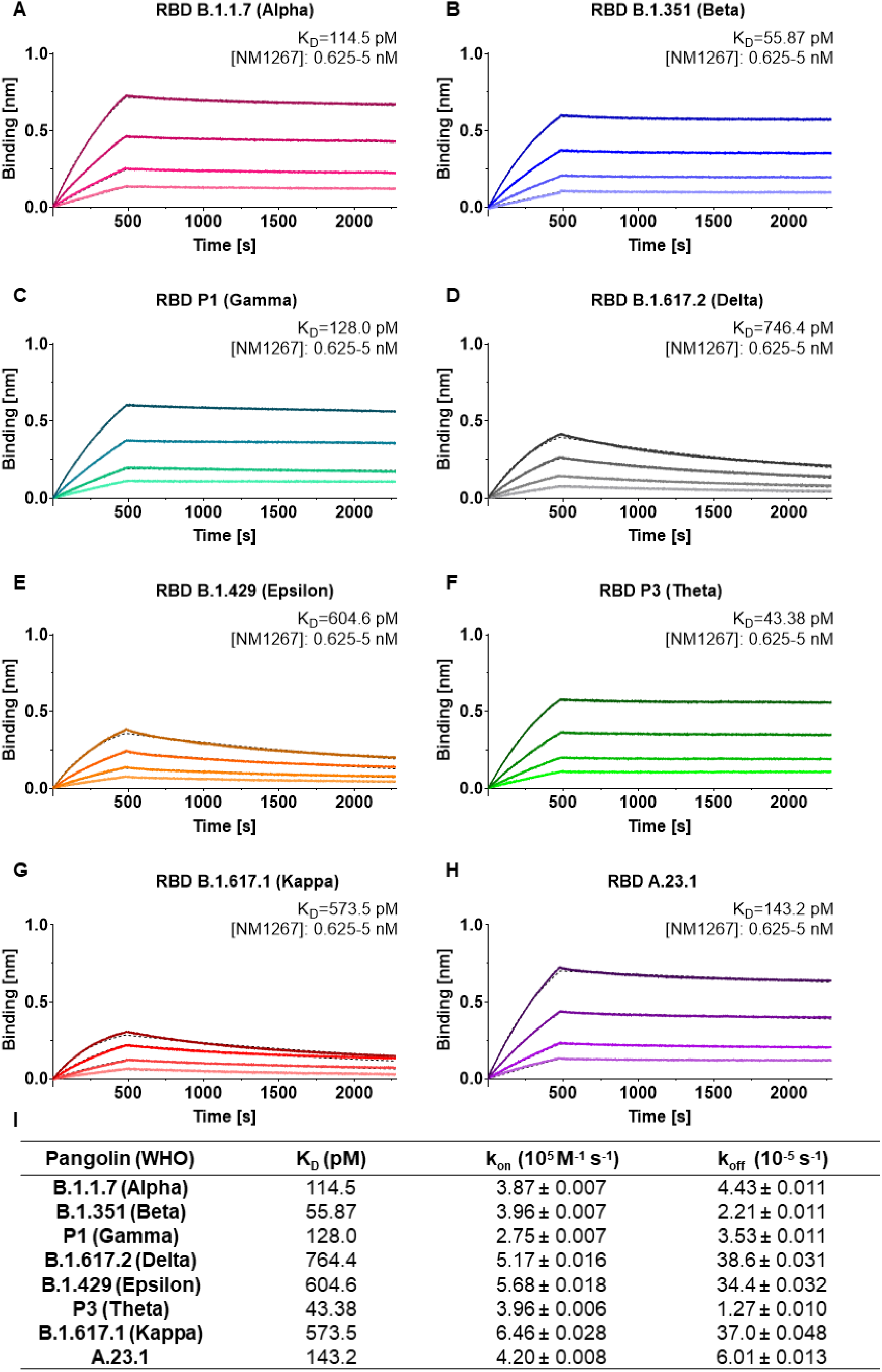
Biparatopic NM1267 targets several recently identified RBD variants with picomolar affinity. **A-H** Affinity measurements by BLI of bipNb NM1267 on recently identified RBD variants B.1.1.7 (Alpha) (**A**), B.1.351 (Beta) (**B**), P1 (Gamma) (**C**), A.1.617.2 (Delta) (**D**), B.1.429 (Epsilon) (**E**), P3 (Theta) (**F**), B.1.617.1 (Kappa) (**G**) and A.23.1 (**H**). NM1267 was applied with concentrations ranging from 5–0.625 nM (illustrated with gradually lighter shades) on immobilized RBD variants. Global 1:1 fits are illustrated as dashed lines. **I** Tabular summary of binding affinity (K_D_), association (k_on_) and dissociation constant (k_off_) determined for the individual RBD variants.

Preceding studies have shown that the B.1.351 (Beta) variant is able to evade immune response after vaccination or treatment with already established Nabs (Madhi *et al.*, 2021; Wang *et al.*, 2021), probably due to the three distinct escape mutations within the RBD (K417N, E484K, N501Y) (Li *et al*, 2021; Zhou *et al.*, 2021). Therefore, determination of the neutralization capacity of NM1267 against a clinical isolate of B.1.351 (Beta) SARS-CoV-2 (Becker *et al.*, 2021) was of particular interest. Performing VNTs, human Caco-2 cells were co-incubated with serial dilutions of NM1267 and WT or B.1.351 (Beta) SARS-CoV-2. As negative control a non-specific bivalent Nb (bivNb) NM1251 was applied in the same setting. Immunofluorescence (IF) staining of the virus was performed 48 h after infection, and infection rates were determined by automated fluorescence microscopy. NM1267 showed strong neutralization of both variants, SARS-CoV-2 WT and SARS-CoV-2 B.1.351 (Beta), with IC_50_ values of 0.33 nM and 0.78 nM, respectively, whereas no effect was observed for the treatment with NM1251 (**Figure 3A-C**).

**Figure 3.**
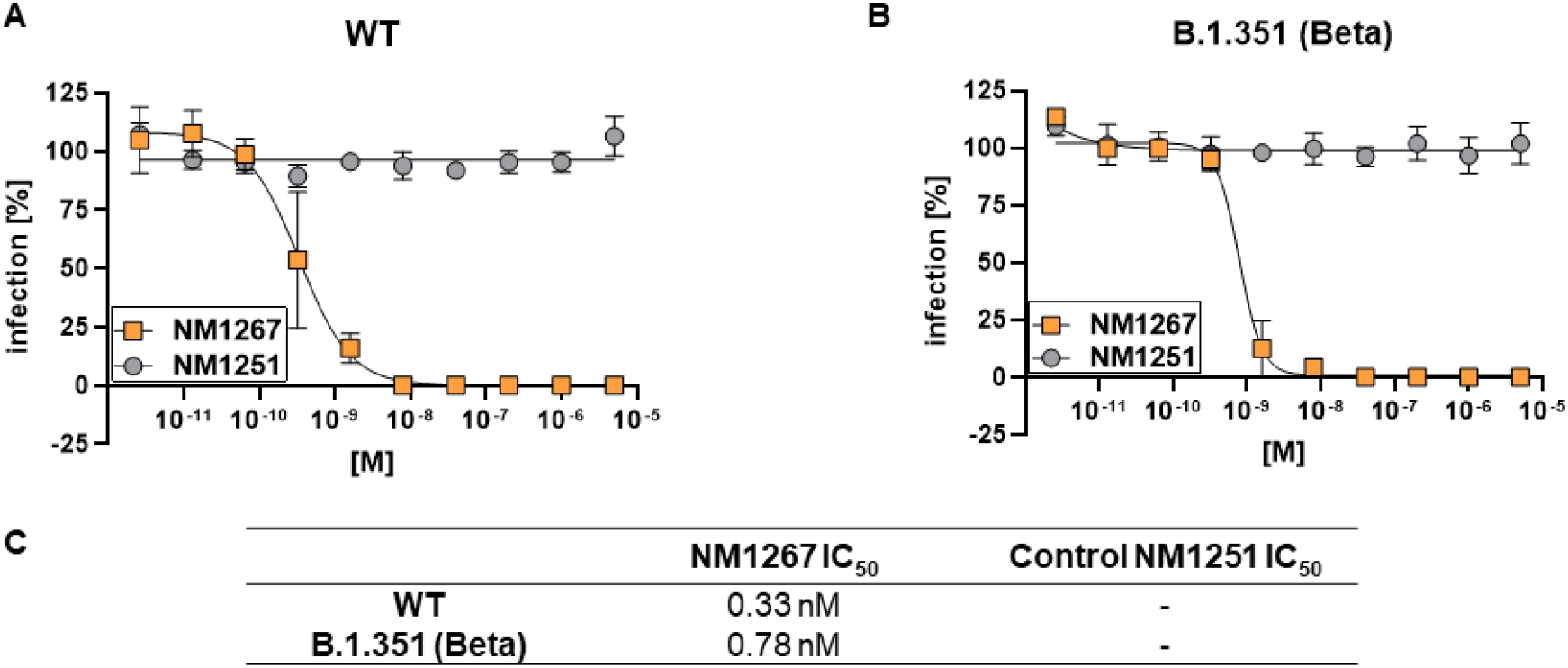
Biparatopic NM1267 neutralizes wild-type and B.1.351 SARS-CoV-2 infection in Caco-2 cells. **A, B** Neutralization potency of NM1267 was analyzed in Caco-2 cells using the SARS-CoV-2 WT (**A**) and SARS-CoV-2 B.1.351 (Beta) (**B**). Infection normalized to virus-only infection control is illustrated as percent of infection (infection [%]). Data are presented as mean ± SEM of three biological replicates (n = 3). **C** Tabular summary of IC_50_ values, calculated from a four-parametric sigmoidal model.

To examine the *in vivo* potency of NM1267, we utilized transgenic K18-hACE2 mice expressing human ACE2, which are highly permissive for infection with human SARS-CoV-2 isolates (Winkler *et al.*, 2020). Considering the broad applicability for which noninvasive routes of administration are preferred, we chose intranasal administration of NM1267. Mice were treated in a prophylactic treatment setting with either 20 μg of NM1267 or the non-specific control (NM1251), followed by SARS-CoV-2 WT or B.1.351 (Beta) infection 7 h later (**Figure 4A**). Weight loss and survival of infected mice were monitored for 14 days post-infection (d p.i.). All WT virus infected animals treated with the negative control NM1251 became severely sick with obvious clinical signs of pneumonia, lost substantial amounts of body weight, and 14 out of 15 animals had to be euthanized (**Figure 4B, C**). In contrast, administration of the SARS-CoV-2 neutralizing NM1267 significantly reduced signs of disease, weight loss and 9 out of 12 animals survived the infection. Additionally, virus shedding by NM1267-treated mice, determined by viral load on nasal swabs, was significantly reduced in comparison with control animals (**Figure D**). Infection of non-specific NM1251-treated mice with the VOC B.1.351 (Beta) also induced substantial weight loss, severe disease symptoms, and 7 out of 9 animals had to be euthanized (**Figure 4E, F**). Similarly to what was observed for the WT virus infection, only one NM1267-treated mouse infected with B.1.351 (Beta) lost substantial amounts of weight and had to be euthanized, whereas 5 out of 6 animals did not show any signs of disease and survived the infection.

**Figure 4.**
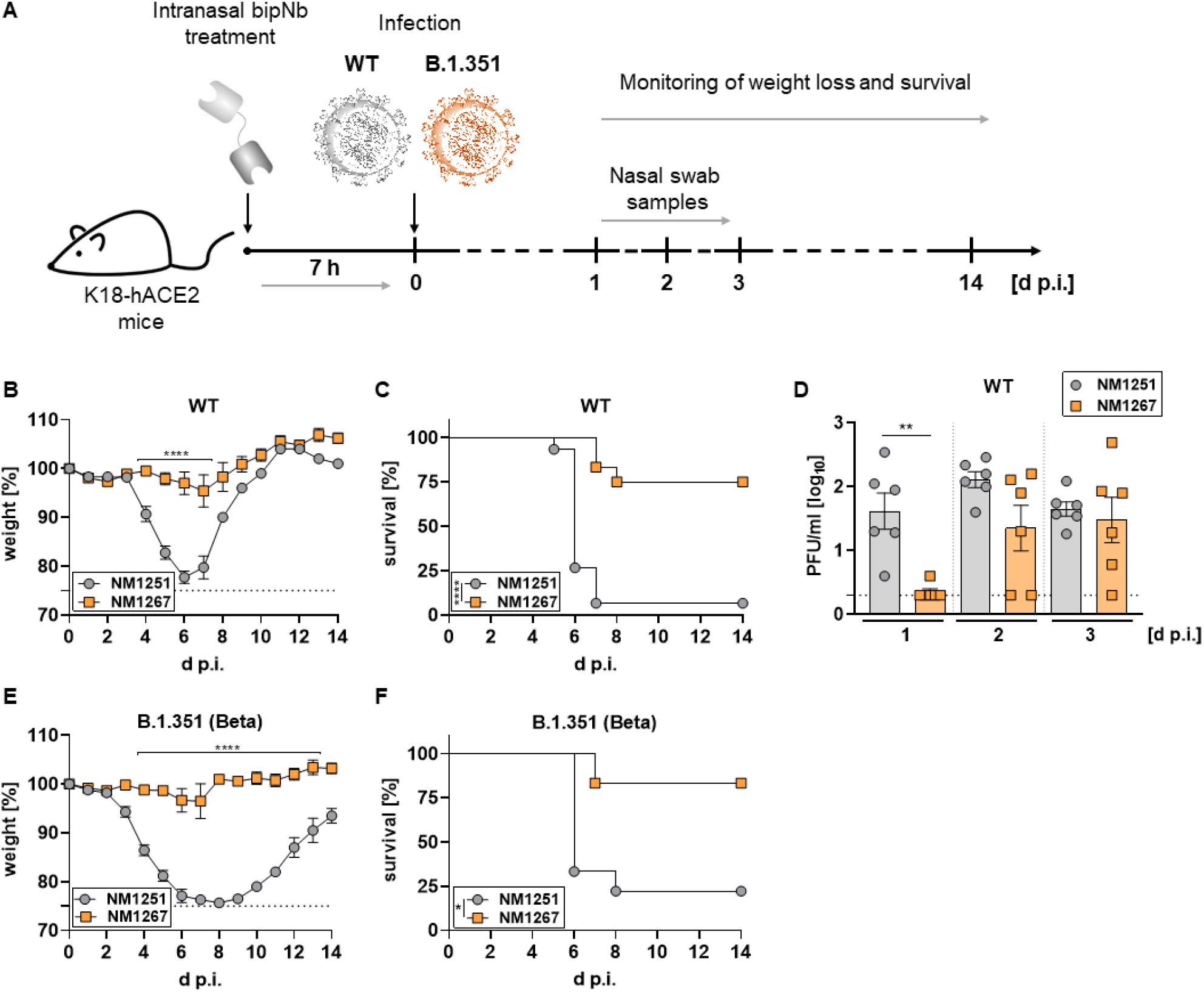
Intranasal application of NM1267 protects K18-hACE2 mice against SARS-CoV-2 induced disease and reduces mortality and virus shedding. **A** Schematic illustration of treatment scheme. **B-D** Hemizygous K18-hACE2 mice were treated intranasally with 20 μg of NM1251 (n = 15) or NM1267 (n = 12) seven hours prior to infection with 3*10^3^ PFU SARS-CoV-2 WT. Weight loss (**B**) and survival (**C**) were monitored for 14 days. Nasal swabs were collected from six mice per group at the indicated time points and viral load determined by plaque-assay (**D**). Symbols represent mean ± SEM in (B) or individual animals in (D). Bars in (D) represent mean ± SEM \*\*\*\**P* < 0.0001, by two-way ANOVA with Sidak’s multiple comparisosns test in (B), \*\*\*\**P* < 0.0001, by log-rank test in (C) and \*\**P* < 0.01, by unpaired t test in (D). **E-F** Hemizygous K18-hACE2 mice were treated intranasally with 20 μg of NM1251 (n = 9) or NM1267 (n = 6) seven hours prior to infection with 3*10^3^ PFU SARS-CoV-2 B.1.351 (Beta). Weight loss (**E**) and survival (**F**) were monitored for 14 days. Symbols in (E) represent mean ± SEM. \*\*\*\**P* < 0.0001, by 2way ANOVA with Sidak’s multiple comparisons test in (E) and \**P* < 0.05, by log-rank test in (F).

Next, we performed histopathological analyses of infected lung tissue samples from treated mice to evaluate the degree of tissue damage upon infection by hematoxylin and eosin (H&E) staining, and the extent and localization of viral RNA-positive lung tissue by *in situ* hybridization (ISH). Applying a grading system from 0 (no tissue damage) to 4 (strong tissue damage), it became evident that all SARS-CoV-2 infected mice under control treatment (NM1251) exhibited a pronounced inflammation and loss of functional lung epithelia (**Figure 5A-C**). In particular, infection with the B.1.351 (Beta) variant caused massive tissue damage in NM1251-treated mice, reaching highest scores between 3 and 4. Notably, prophylactic treatment with NM1267 efficiently reduced virus- and inflammation-induced tissue damage within the lungs (scoring 0.5-1.5) of both, SARS-CoV-2 WT and B.1.351 (Beta) infected mice (**Figure 5A-C**). In line with these findings, distinctly lower levels of SARS-CoV-2 RNA were found in samples taken from NM1267-treated mice, restricted to small areas of the lung at sub-pleural position and some fat cells. In contrast, tissue sections of control-treated (NM1251) mice showed that epithelial cells in large areas of the lung were virus RNA positive, independent of the SARS-CoV-2 variant (**Figure 5B**). In summary, these data provide strong evidence that intranasal application of bipNb NM1267 successfully prevents SARS-CoV-2-induced tissue damage, disease progression, and that the prophylactic treatment reduces virus shedding and mortality *in vivo*.

**Figure 5.**
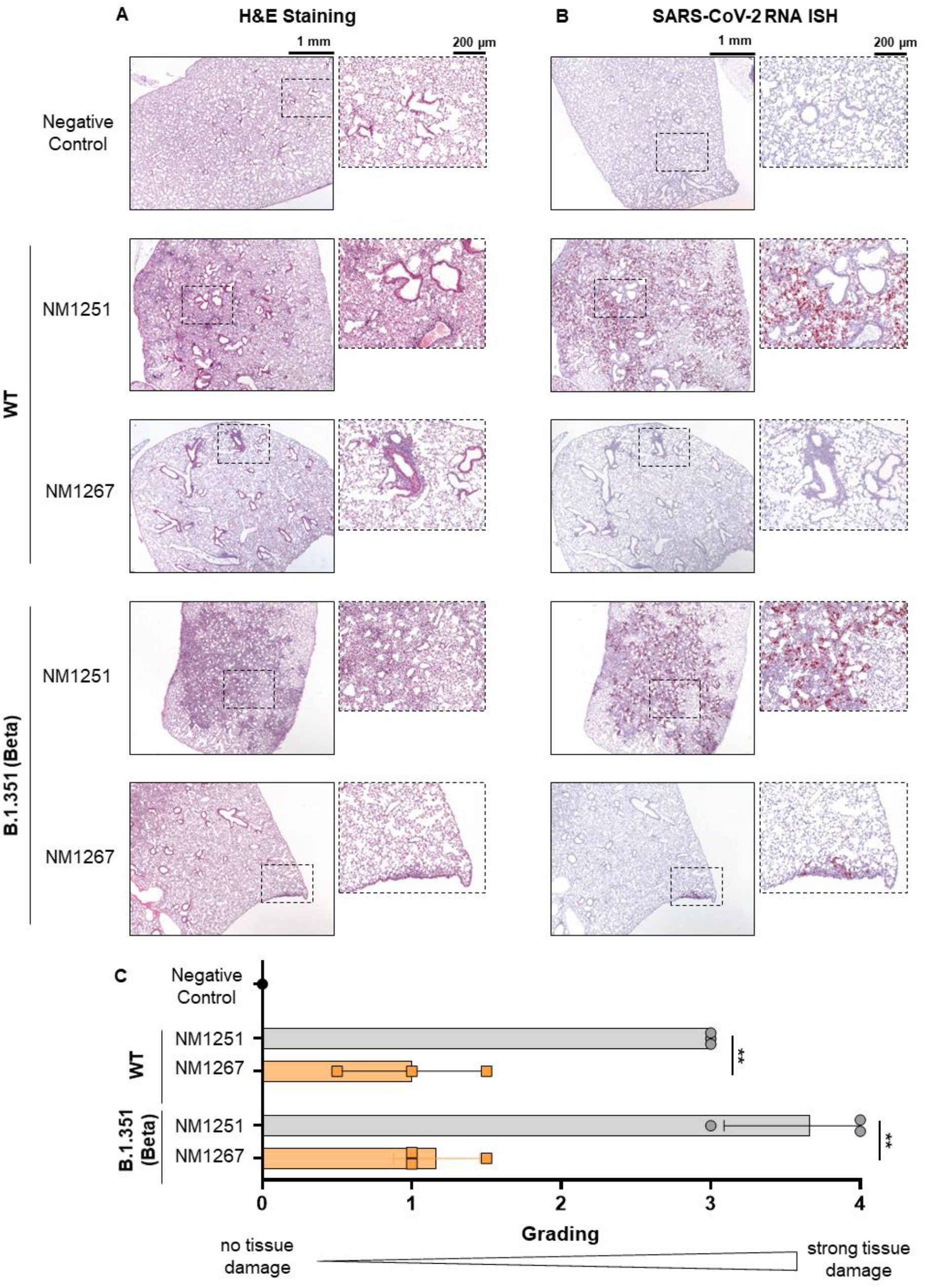
Microscopic analysis of lung tissue from SARS-CoV-2 infected K18-hACE2 mice. Mice were intranasally treated with bipNB NM1267 or control Nb NM1251 and subsequently infected with 3*10^3^ PFU SARS-CoV-2 WT or B.1.351 (Beta) variant. **A-B** Serial tissue sections revealed severe inflammation (H&E) and numerous widespread SARS-CoV-2 RNA positive alveolar epithelia cells and macrophages (*in situ* hybridization (ISH)) in lungs of infected, control-treated mice. In infected and NM1267 bipNb treated animals no inflammation or only small focal areas with inflammation and a few SARS-CoV-2 RNA positive cells were observed. **C** Quantification of lung damage was done in n = 3 animals per group and grading score of individual animals is presented as mean ± SD with \*\**P* < 0.01, by unpaired t test.

## Discussion

The ongoing COVID-19 pandemic and the frequent emergence of new SARS-CoV-2 variants of concern highlight the need of easily applicable therapeutic options. In addition to Nabs, Nbs targeting the RBD of SARS-CoV-2 offer a promising alternative. Not only their stable folding, robust biochemical properties, and ease of functionalization/ multimerization, but also the lack of an Fc moiety that prevents severe adverse events such as ADE and their low immunogenicity underlines the potential of Nbs as advanced therapeutic tools (Muyldermans, 2013; Taylor *et al*, 2015; Tirado & Yoon, 2003). In this context, prophylactic or therapeutic administration of neutralizing Nbs have already been shown not only to limit virus replication and weight loss in animals, but also to minimize lung damage and mortality in transgenic hACE2 mice and Syrian hamsters after infection with SARS-CoV-2 variants (Haga *et al*, 2021; Hanke *et al*, 2021; Nambulli *et al.*, 2021; Schepens *et al.*, 2021; Wrapp *et al.*, 2020).

Recently, we identified neutralizing Nbs and showed their applicability to quantify neutralizing antibodies in serum from convalescent and immunized individuals (Becker *et al.*, 2021; Wagner *et al.*, 2021). In this study, we investigated the therapeutic potential of multiple biparatopic Nbs that jointly target different epitopes within the RBD. All bipNbs revealed comparable picomolar affinities to RBD_WT_, strong ACE2 displacement and high neutralization capacities. However, since NM1267 showed the highest thermal stability and detailed data on the recognized epitopes were already available (Wagner *et al.*, 2021), we selected NM1267 as most promising drug candidate. Hypothesizing that combinatorial binding to epitopes within the RBD:ACE2 interface and to conserved epitopes outside this interaction site might be beneficial to also cover emerging VOCs, we were subsequently able to show strong binding of NM1267 to all SARS-CoV-2 RBD variants tested. We further, demonstrated a high neutralizing capacity of NM1267 for SARS-CoV-2 WT and the B.1.351 (Beta) variant in VNTs. Use of NM1267 for prophylactic intranasal application strongly diminished disease progression in both SARS-CoV-2 WT and B.1.351 (Beta) infected mice and resulted in 4.6- to 11.2-fold increased survival rates compared to control-treated animals. Overall, these data underscore the potential of NM1267 to treat infections with VOCs for which currently available vaccines and therapeutic approaches are suspected to have reduced efficacy (Becker *et al.*, 2021; Kustin *et al.*, 2021; Madhi *et al.*, 2021; Planas *et al*, 2021; Zhou *et al.*, 2021). Histopathological analyses and *in situ* hybridization detecting viral RNA in lung tissue samples further revealed dramatically reduced tissue damage and viral load suggesting that NM1267 treatment may also reduce long-term effects of SARS-CoV-2 infections (Han *et al*, 2021; Yong, 2021).

Notably, most strategies for engineering neutralizing Nbs currently rely on increasing avidity by generating multivalent constructs binding the same epitope (Nambulli *et al.*, 2021; Schepens *et al.*, 2021; Wu *et al*, 2021). To date, solely *Hanke et al.* followed a similar strategy as suggested in this study, by coupling two different Nbs, Fu2 and Ty1, to generate a potential therapeutic molecule. However, these two Nbs recognize overlapping epitopes at the RBD:ACE2 interaction site (Hanke *et al.*, 2021), which could facilitate virus escape.

The administration of NM1267 showed strong short-term efficacy *in vivo*. However, certain modifications like fusion to an Fc-moiety, albumin binding motif or directly to carrier proteins like albumin may improve duration of effectiveness (Hanke *et al.*, 2021; Nambulli *et al.*, 2021; Schepens *et al.*, 2021; Wu *et al.*, 2021). Moreover, the extreme susceptibility of transgenic hACE2 mice to SARS-CoV-2-induced disease due to the artificial overexpression of hACE2 in a variety of tissues and organs, may even result in an underestimated therapeutic potential of NM1267. To address this issue and to investigate also the potential of NM1267 to prevent active transmission of SARS-CoV-2 via direct contact and aerosol, further studies in more physiological models, such as Syrian hamsters or non-human primates, would be required (Haga *et al.*, 2021; Nambulli *et al.*, 2021; Schepens *et al.*, 2021).

In summary, with the development and detailed characterization of the neutralizing potential of NM1267 in combination with intranasal *in vivo* application, we offer a straightforward prophylactic, and possibly therapeutic, approach to combat infections with emerging VOCs. Given the poor access to vaccines in various countries, vaccination fatigue and the frequent emergence of new variants of concern, we believe that the development of such easily applicable therapeutic approaches to protect and treat predisposed individuals are highly promising strategies and urgently warranted.

## Material & Methods

### Expression constructs

To generate described expression constructs all used primer sequences are listed in **Supplementary Table 2**. Nb NM1267 was generated as described previously (Wagner *et al.*, 2021). To generate bipNb NM1266, Nb NM1230 was genetically fused via internal (G_4_S)_4_– linker to the N-terminus of Nb NM1228 (Wagner *et al.*, 2021). Nb cDNAs were PCR amplified by the use of primers NM1230Nfor, NM1230Nrev and NM1228Cfor, NM1228Nrev and subsequently fused by overlap extension PCR. BipNb NM1268 composed of Nb NM1228 and Nb NM1226 (Wagner *et al.*, 2021) was similarly generated using primers NM1228Nfor, NM128Nrev and NM1226Cfor, NM1226Crev. DNA coding for bipNbs were cloned into pCDNA3.4 expression vector seamlessly downstream of comprising N-terminal signal peptide (MGWTLVFLFLLSVTAGVHS) for secretory pathway using type IIS restriction enzyme Esp3I and EcoRI site. Coding sequence of bivNb NM1251 (Traenkle *et al*, 2020) was produced by gene synthesis (Thermo Fisher Scientific, Massachusetts, USA) and similarly cloned into pCDNA3.4 expression vector.

Receptor binding domain (RBD) variants of SARS-CoV-2 were generated as earlier published (Wagner *et al.*, 2021). The expression plasmid pCAGGS encoding the receptor-binding domain (RBD) of SARS-CoV-2 spike protein (amino acids 319-541) was kindly provided by F. Krammer. RBDs of SARS-CoV-2 variants of concern (VOCs) B.1.1.7 (Alpha), B.1.351 (Beta), P1 (Gamma), A.1.617.2 (Delta), B.1.429 (Epsilon), P3 (Theta), B.1.617.1 (Kappa) and A.23.1 were generated by PCR amplification of fragments from WT or cognate DNA template and subsequent fusion PCR by overlap extension to introduce described mutations. Based on RBD_WT_ sequence primer pairs RBDfor N501Yrev and N501Yfor RBDrev were used for the amplification of B.1.1.7 (Alpha) sequence; primer pairs RBDfor L452Rrev and L452Rfor RBDrev for B.1.429 (Epsilon); RBDfor, V367Frev and V367Ffor, RBDrev for A.23.1. A.1.617.2 (Delta) was generated based on B.1.429 (Epsilon) using primer pairs RBDfor T478Krev and T478Kfor RBDrev. Based on B.1.1.7 (Alpha) sequence P3 (Theta) was generated using primer pairs RBDfor E484Krev and E484Kfor RBDrev. B.1.617.1 (Kappa) was generated using primer pairs RBDfor E484Krev and E484Kfor RBDrev as well as RBDfor L452Rrev and L452Rfor RBDrev. B.1.351 (Beta) and P1 (Gamma) were generated based on P3 (Theta) sequence using primer pairs RBDfor K417Nrev and K417Nfor RBDrev; and RBDfor K417Trev and K417Tfor RBDrev, respectively.

Amplicons were inserted using XbaI and NotI site into the pCDNA3.4 expression vector. The integrity of all expression constructs was confirmed by standard sequencing analysis.

### Protein expression and purification

Confirmed constructs were expressed in Expi293 cells. Briefly, cells were cultivated (37°C, 125 rpm, 8% (v/v) CO_2_) to a density of 5.5 × 10^6^ cells/ mL, diluted with Expi293F expression medium and transfection of the corresponding plasmids (1 μg/mL) with expifectamine. 20 h post transfection enhancers were added as per the manufacturer’s instructions. Cell suspensions were then cultivated for 2–5 days (37 °C, 125 rpm, 8% (v/v) CO_2_) and then centrifuged (4°C, 23,900×*g*, 20 min) to clarify the supernatant. Supernatants were then filtered with a 0.22-μm membrane (Millipore, Darmstadt, Germany) and supplemented with His-A buffer stock solution (20 mM Na_2_HPO_4_, 30 mM NaCl, 20 mM imidazole, pH 7.4). The solution was then applied to a HisTrap FF crude column on an Aekta pure system (GE Healthcare, Freiburg, Germany), extensively washed with His-A buffer, and eluted with an imidazole gradient (50–400 mM). Buffer exchange to PBS and concentration of eluted proteins were carried out using Amicon 10 K centrifugal filter units (Millipore, Darmstadt, Germany). Protein quality was analyzed by standard SDS-Page and via the NanoDrop protein concentration was determined.

### Affinity measurements

Binding affinity of bipNbs towards variants of RBD was determined via biolayer interferometry (BLI) using the Octet RED96e system according to standard protocol. Therefore, RBD variants were biotinylated and immobilized on streptavidin biosensors (SA, Sartorius). Dilution series ranging from 5-0.625 nM of bipNbs were applied and one reference was included per run. For affinity determination, the 1:1 global fit of the Data Analysis HT 12.0 software was used.

### Bead-based multiplex ACE2 competition assay

To analyze binding competition of human ACE2 versus generated bipNbs the bead-based multiplex ACE2 competition assay was performed as previously described (Wagner *et al.*, 2021).

### Stability analysis

Stability analysis was performed by the Prometheus NT.48 (Nanotemper). Therefore, freshly-thawed bipNbs were diluted to 0.25 mg/mL and measurements were carried out at time point T_0_ and after incubation for ten days at 37°C (T_10_) using high-sensitivity capillaries. Thermal unfolding and aggregation of the bipNbs is induced by the application of a thermal ramp of 20-95°C, while measuring fluorescence ratios (F350/F330) and light scattering. Via the PR. ThermControl v2.0.4 the melting (T_m_) and aggregation (T_Agg_/ T_turbidity_) temperature was determined.

### Viruses

All experiments associated with the SARS-CoV-2 virus were conducted in Biosafety Level 3 laboratory. The recombinant infectious SARS-CoV-2 clone expressing mNeonGreen (icSARS-CoV-2-mNG) (PMID: 32289263) was obtained from the World Reference Center for Emerging Viruses and Arboviruses (WRCEVA) at the UTMB (University of Texas Medical Branch) (Xie *et al*, 2020) and used as described (Ruetalo *et al*, 2021). SARS-CoV-2 WT (SARS-CoV-2 Tü1 or SARS-CoV-2 Muc IMB-1) and SARS-CoV-2 B.1.351 (Beta) (SARS-CoV SAv) were isolated from patient samples and variant identity was confirmed by next-generation sequencing of the entire viral genome as described in (Ruetalo *et al.*, 2021) and (Becker *et al.*, 2021), respectively.

### Virus neutralization assay (VNT)

Caco-2 (Human Colorectal adenocarcinoma, ATCC HTB-37) cells were cultured at 37°C with 5% CO_2_ in DMEM containing 10% FCS, 2 mM l-glutamine, 100 μg/ ml penicillin-streptomycin and 1% NEAA.

Neutralization assays using clinical isolates (**Figure 3A-C**) were performed as described in (Becker *et al.*, 2021; Ruetalo *et al.*, 2021). Briefly, cells were co-incubated with the respective clinical isolate SARS-CoV-2 WT (200325_Tü1) at an MOI of 0.8 or SARS-CoV-2 B.1.351 (Beta) (210211_SAv) at an MOI of 0.6 and serial dilutions of the bipNb from 5 uM to 0.064 nM. 48 h post-infection, cells were fixed with 80% acetone, and immune fluorescence (IF) staining was performed using an anti-SARS-CoV-2 nucleocapsid antibody (rabbit) and a goat anti-rabbit Alexa594 conjugated secondary antibody. Cells were counterstained with DAPI solution and images were taken with the Cytation3 (BioTek). Infection rates were calculated as the ratio of DAPI-positive over Alexa594-positive cells, which were automatically counted by the Gen5 software (BioTek). In the case of the neutralization assay using the icSARS-CoV-2-mNG (**Supplementary Figure 1A-C**) the protocol was previously described in (Ruetalo *et al.*, 2021; Wagner *et al.*, 2021). Data were normalized to respective virus-only infection control. Inhibitory concentration 50 (IC_50_) was calculated as the half-maximal inhibitory dose using 4-parameter nonlinear regression (GraphPad Prism).

### *In vivo* infection experiments

Transgenic (K18-hACE2)2Prlmn mice were purchased from The Jackson Laboratory and bred and kept under specific-pathogen-free conditions in the animal facilities of the University Medical Center Freiburg. Hemizygous 8-14 week-old animals of both sexes were used in accordance with the guidelines of the Federation for Laboratory Animal Science Associations and the National Animal Welfare Body. All experiments were in compliance with the German animal protection law and approved by the animal welfare committee of the Regierungspraesidium Freiburg (permit G-20/91). Mice were anesthetized using isoflurane and treated intranasally (i.n.) with 20 μg of NM1251 or NM1267 seven hours prior to infection with 3*10^3^ PFU of the respective SARS-CoV-2 isolate (SARS-CoV-2 WT (SARS-CoV-2 Muc-IMB-1) and SARS-CoV B.1.351 (Beta) (SARS-CoV-2 SAv)) in 40 μl PBS containing 0.1% BSA. Infected mice were monitored for weight loss and clinical signs of disease for 14 days and sacrificed if severe symptoms were observed or body weight loss exceeded 25% of the initial weight. Superficial nasal swabs were taken on days 1, 2 and 3 post infection. Swabs were collected in OptiMEM containing 0.3% BSA and titers determined by plaque assay using Vero E6 cells. Infected ketamine/ xylazine-anesthetized mice were prepared for histological analyses by transcardial perfusion with 15 ml of 4% formaldehyde solution and stored in 4 % formaldehyde at 4 °C until organs were processed further. All experiments were performed under BSL3 conditions.

### Haematoxylin and eosin (H&E) staining and *in situ* hybridization (ISH)

Lung tissue was routinely embedded in paraffin and H&E staining was performed from 4 μm thick lung tissue sections by using the Tissue-Tek® Prisma (Sakura, Umkirch, Germany). To detect SARS-CoV-2 RNA (plus-strand RNA), 4 μm thick lung tissue sections, including negative and positive controls, were hybridized using specific probes for SARS-CoV-2 (Advanced Cell Diagnostics (ACD), Newark, CA, USA) followed by the RNAscope 2.5 HD Detection Kit Red from ACD (Newark, CA, USA) according to the manufacturer’s protocol. Quantification of tissue damage including inflammation was defined as grade 0: no damage, grade 1: 1-10%, grade 2: 10-20%, grade 3: 20-50%, grade 4: 50-80% of lung tissue was involved.

## Analyses and Statistics

Graph preparation and statistical analysis was performed using the GraphPad Prism Software (Version 9.0.0 or higher).

## Data availability

The data that support the findings of this study are available from the corresponding authors upon reasonable request.

## Acknowledgements

This work was supported by the Initiative and Networking Fund of the Helmholtz Association of German Research Centers (grant number SO-96), the European Union’s Horizon 2020 research and innovation program under grant agreement No 101003480—CORESMA. This work has further received funding from the State Ministry of Baden-Württemberg for Economic Affairs, Labour and Housing Construction (FKZ 3-4332.62-NMI/68), from the Ministry of Science, Research and Arts of the State of Baden-Württemberg (COVID-19 funding) and from the Deutsche Herzstiftung. This work was supported by the Bundesministerium fuer Bildung und Forschung (BMBF) through the Deutsches Zentrum fuer Luft- und Raumfahrt, Germany, (DLR, grant number 01KI2077) and by the Federal State of Baden-Wuerttemberg, Germany, MWK-Sonderfoerdermaßnahme COVID-19/AZ.:33-7533.-6-21/7/2 to MS. We thank Florian Krammer for providing expression constructs for SARS-CoV-2 homotrimeric Spike and RBD.

## Authorship Contributions

Study design: TRW, DS, MiS, MS, UR; Nb biochemical characterization: TRW, PDK, BT, DIF; Multiplex binding assay: DJ, MB, NSM; Virus neutralization assays: NR, MiS; mouse infection experiments: DS, JB, AO; histopathological analysis and *in situ* hybridization: KK, MSa; Data analysis and statistical analysis: TRW, DS, JB, NR, MiS, KK, MS, UR; Manuscript drafting: TRW, UR; Study supervision: MS, UR; Manuscript reviewing and editing: All authors.

## Supplementary Information

**Supplementary Figure 1.**
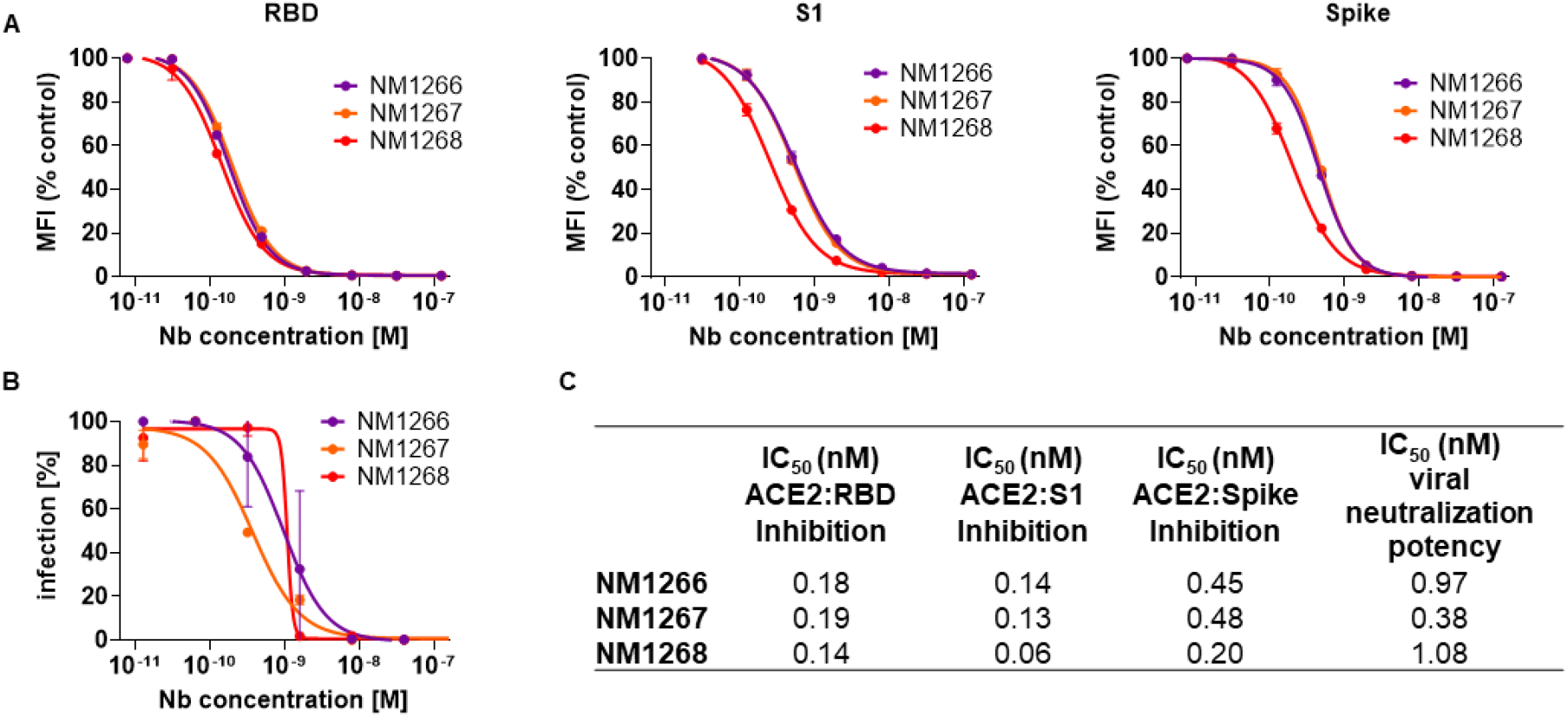
Biparatopic NM1266, NM1267 and NM1268 compete with ACE2 and neutralizes SARS-CoV-2 infection. **A** Results from multiplex ACE2 competition assay are shown for the three spike-derived antigens: RBD, S1-domain (S1), and homotrimeric spike (Spike). Color-coded beads coated with the respective antigens were co-incubated with biotinylated ACE2 and dilution series of NM1266, NM1267 and NM1268 (8 pM to 126 nM) followed by measuring residual binding of ACE2. MFI signals were normalized to the maximum detectable signal per antigen given by the ACE2-only control. IC_50_ values were calculated from a four-parametric sigmoidal model. Data are presented as mean ± SD of three technical replicates. **B** Neutralization potency of NM1266, NM1267 and NM1268 was analyzed in Caco-2 cells using the SARS-CoV-2-mNG infectious clone. Infection rate normalized to virus-only infection control is illustrated as percent of infection (infection [%]). IC_50_ value was calculated from a four-parametric sigmoidal model, and data are presented as mean ± SEM of three biological replicates (n = 3). **C** Table summarizing IC_50_ values of the multiplex ACE2 competition assay and virus neutralization assay obtained for NM1266, NM1267 and NM1268.

**Supplementary Figure 2.**
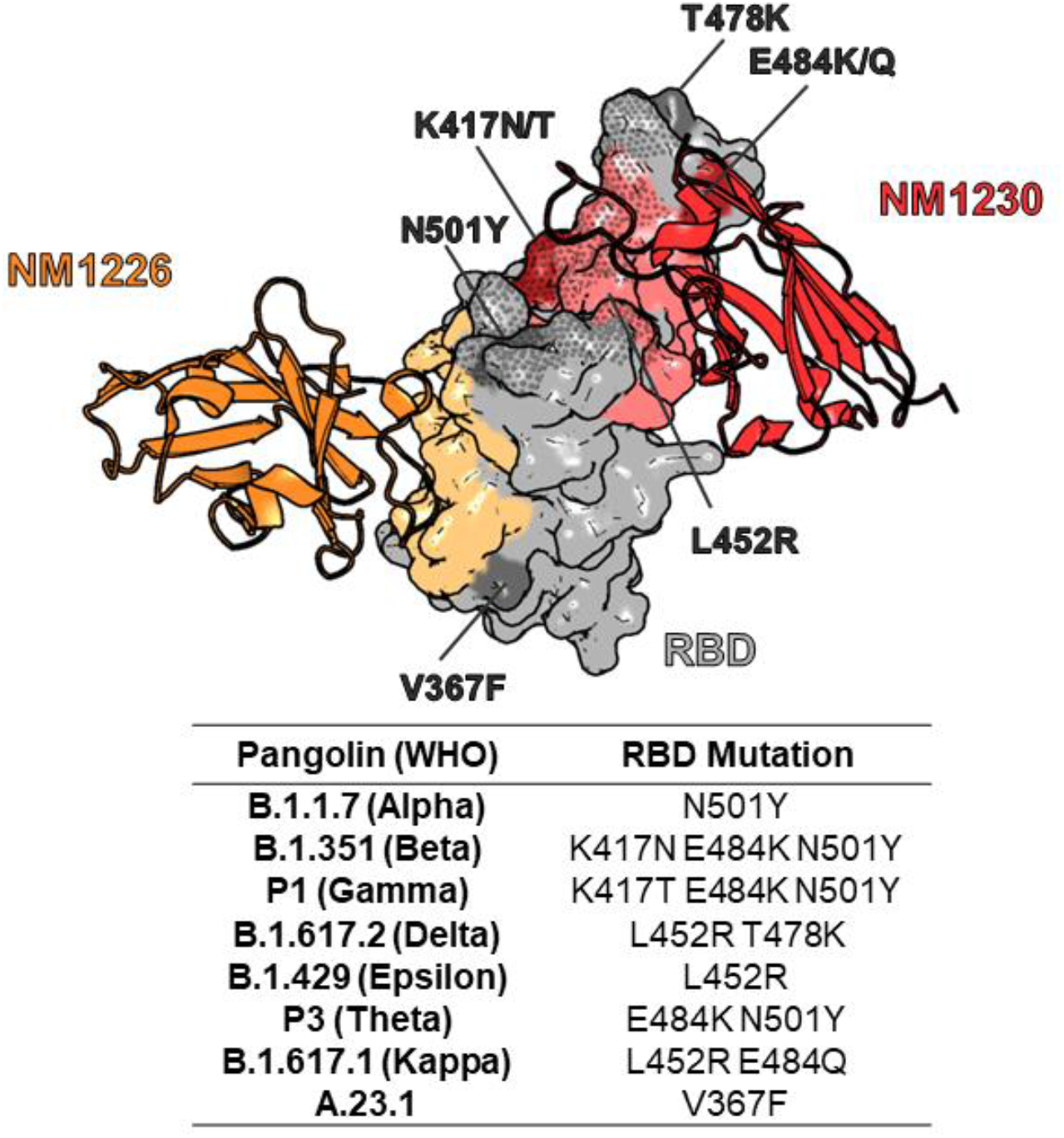
Influence of RBD mutations on bipNb NM1267 binding. NM1267-froming single Nbs, NM1226 (orange, PDB 7NKT) and NM1230 (red, PDB 7B27) are shown as cartoon with their corresponding binding epitopes on the RBD surface in light orange and light red, respectively. In addition, the ACE2 interaction site on RBD is illustrated as dotted surface. Mutations on RBD of identified SARS-CoV-2 variants, including B.1.1.7 (Alpha), B.1.351 (Beta), P1 (Gamma), A.1.617.2 (Delta), B.1.429 (Epsilon), P3 (Theta), B.1.617.1 (Kappa) and A.23.1 are highlighted in dark grey or dark red and labeled respectively.

**Supplementary Table 1.** Nb combinations for bipNbs.

| Biparatopic (Bip)/ bivalent Nb | Single Nb combination |
| --- | --- |
| NM1266 | NM1230-(G<sub>4</sub>S)<sub>4</sub>-NM1228 |
| NM1267 | NM1230-(G<sub>4</sub>S)<sub>4</sub>-NM1226 |
| NM1268 | NM1228-(G<sub>4</sub>S)<sub>4</sub>-NM1226 |
| NM1251 (Control) | Pep-Nb-(G<sub>4</sub>S)<sub>4</sub>-Pep-Nb |

**Supplementary Table 2.**
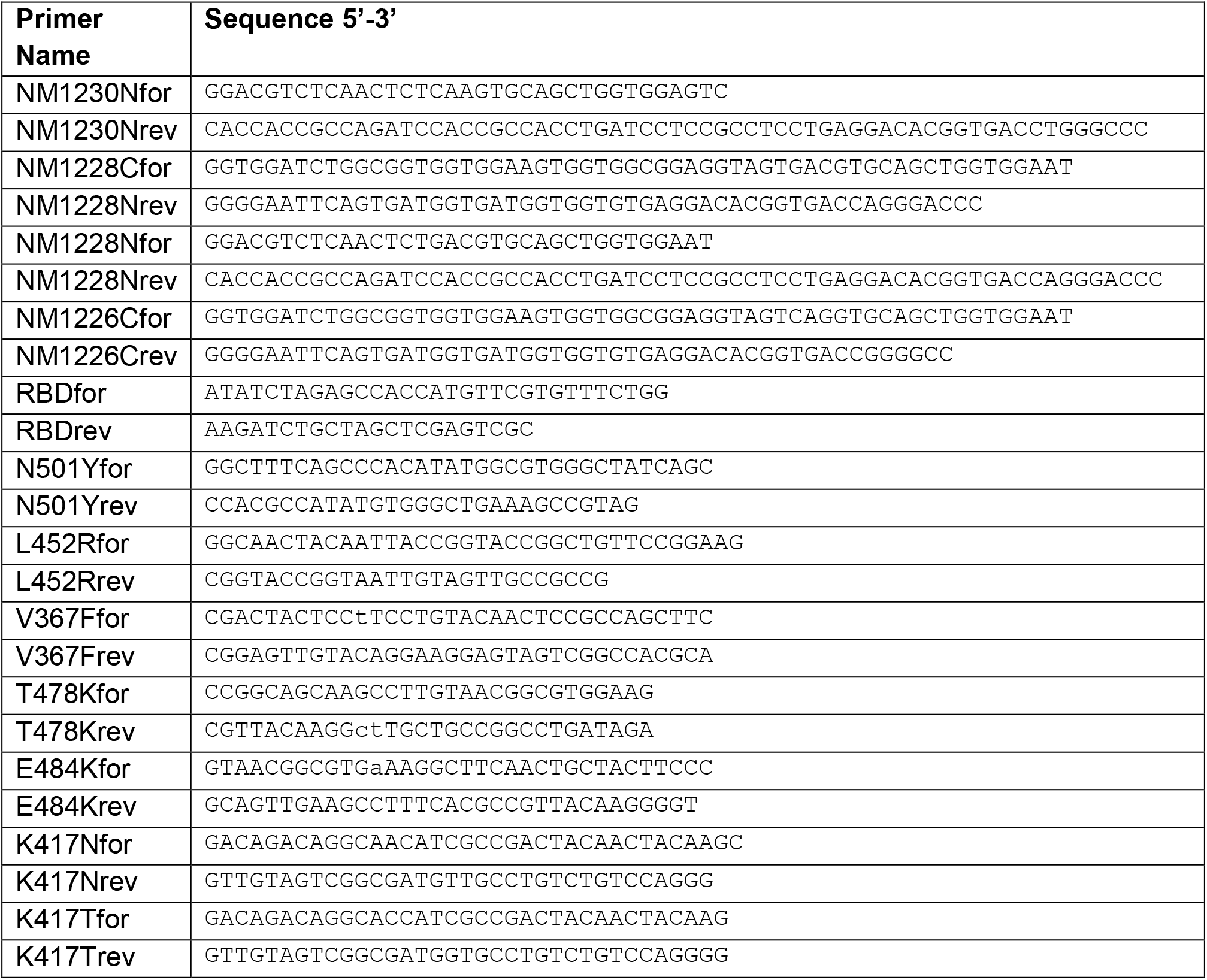
Primer Sequences.

## References

Beaudoin-Bussières G, Laumaea A, Anand SP, Prévost J, Gasser R, Goyette G, Medjahed H, Perreault J, Tremblay T, Lewin A (2020) Decline of humoral responses against SARS-CoV-2 spike in convalescent individuals. MBio 11

Becker M, Dulovic A, Junker D, Ruetalo N, Kaiser PD, Pinilla YT, Heinzel C, Haering J, Traenkle B, Wagner TR et al (2021) Immune response to SARS-CoV-2 variants of concern in vaccinated individuals. Nature Communications 12: 3109

Brouwer PJ, Caniels TG, van der Straten K, Snitselaar JL, Aldon Y, Bangaru S, Torres JL, Okba NM, Claireaux M, Kerster G (2020) Potent neutralizing antibodies from COVID-19 patients define multiple targets of vulnerability. Science 369: 643–650

Cao Y, Su B, Guo X, Sun W, Deng Y, Bao L, Zhu Q, Zhang X, Zheng Y, Geng C (2020) Potent neutralizing antibodies against SARS-CoV-2 identified by high-throughput single-cell sequencing of convalescent patients’ B cells. Cell 182: 73–84. e16

Challen R, Brooks-Pollock E, Read JM, Dyson L, Tsaneva-Atanasova K, Danon L (2021) Risk of mortality in patients infected with SARS-CoV-2 variant of concern 202012/1: matched cohort study. bmj 372

Chen P, Nirula A, Heller B, Gottlieb RL, Boscia J, Morris J, Huhn G, Cardona J, Mocherla B, Stosor V et al (2020) SARS-CoV-2 Neutralizing Antibody LY-CoV555 in Outpatients with Covid-19. New England Journal of Medicine 384: 229–237

Chi X, Liu X, Wang C, Zhang X, Li X, Hou J, Ren L, Jin Q, Wang J, Yang W (2020) Humanized single domain antibodies neutralize SARS-CoV-2 by targeting the spike receptor binding domain. Nature communications 11: 1–7

Dagan N, Barda N, Kepten E, Miron O, Perchik S, Katz MA, Hernán MA, Lipsitch M, Reis B, Balicer RD (2021) BNT162b2 mRNA Covid-19 vaccine in a nationwide mass vaccination setting. New England Journal of Medicine 384: 1412–1423

Davies NG, Abbott S, Barnard RC, Jarvis CI, Kucharski AJ, Munday JD, Pearson CAB, Russell TW, Tully DC, Washburne AD et al (2021a) Estimated transmissibility and impact of SARS-CoV-2 lineage B.1.1.7 in England. Science 372: eabg3055

Davies NG, Jarvis CI, van Zandvoort K, Clifford S, Sun FY, Funk S, Medley G, Jafari Y, Meakin SR, Lowe R et al (2021b) Increased mortality in community-tested cases of SARS-CoV-2 lineage B.1.1.7. Nature

Diamond M, Chen R, Xie X, Case J, Zhang X, VanBlargan L, Liu Y, Liu J, Errico J, Winkler E (2021) SARS-CoV-2 variants show resistance to neutralization by many monoclonal and serum-derived polyclonal antibodies. Research square: rs. 3. rs-228079

Haga K, Takai-Todaka R, Matsumura Y, Takano T, Tojo T, Nagami A, Ishida Y, Masaki H, Tsuchiya M, Ebisudani T et al (2021) Nasal delivery of single-domain antibodies improves symptoms of SARS-CoV-2 infection in an animal model. bioRxiv: 2021.2004.2009.439147

Han X, Fan Y, Alwalid O, Li N, Jia X, Yuan M, Li Y, Cao Y, Gu J, Wu H (2021) Six-month follow-up chest CT findings after severe COVID-19 pneumonia. Radiology 299: E177–E186

Hanke L, Das H, Sheward DJ, Vidakovics LP, Urgard E, Moliner-Morro A, Karl V, Pankow A, Kim C, Smith NL et al (2021) A bispecific monomeric nanobody induces SARS-COV-2 spike trimer dimers. bioRxiv: 2021.2003.2020.436243

Hanke L, Perez LV, Sheward DJ, Das H, Schulte T, Moliner-Morro A, Corcoran M, Achour A, Hedestam GBK, Hällberg BM (2020) An alpaca nanobody neutralizes SARS-CoV-2 by blocking receptor interaction. Nature communications 11: 1–9

Huo J, Le Bas A, Ruza RR, Duyvesteyn HM, Mikolajek H, Malinauskas T, Tan TK, Rijal P, Dumoux M, Ward PN (2020) Neutralizing nanobodies bind SARS-CoV-2 spike RBD and block interaction with ACE2. Nature structural & molecular biology 27: 846–854

Jewell BL (2021) Monitoring differences between the SARS-CoV-2 B. 1.1. 7 variant and other lineages. The Lancet Public Health

Jiang S, Hillyer C, Du L (2020) Neutralizing antibodies against SARS-CoV-2 and other human coronaviruses. Trends in immunology 41: 355–359

Ju B, Zhang Q, Ge J, Wang R, Sun J, Ge X, Yu J, Shan S, Zhou B, Song S (2020) Human neutralizing antibodies elicited by SARS-CoV-2 infection. Nature 584: 115–119

Koenig P-A, Das H, Liu H, Kümmerer BM, Gohr FN, Jenster L-M, Schiffelers LD, Tesfamariam YM, Uchima M, Wuerth JD (2021) Structure-guided multivalent nanobodies block SARS-CoV- 2 infection and suppress mutational escape. Science 371

Kustin T, Harel N, Finkel U, Perchik S, Harari S, Tahor M, Caspi I, Levy R, Leshchinsky M, Dror SK (2021) Evidence for increased breakthrough rates of SARS-CoV-2 variants of concern in BNT162b2-mRNA-vaccinated individuals. Nature Medicine: 1–6

Kwok KO, Lai F, Wei WI, Wong SYS, Tang JW (2020) Herd immunity–estimating the level required to halt the COVID-19 epidemics in affected countries. Journal of Infection 80: e32–e33

Li Q, Nie J, Wu J, Zhang L, Ding R, Wang H, Zhang Y, Li T, Liu S, Zhang M (2021) SARS- CoV-2 501Y. V2 variants lack higher infectivity but do have immune escape. Cell 184: 2362–2371. e2369

Long Q-X, Tang X-J, Shi Q-L, Li Q, Deng H-J, Yuan J, Hu J-L, Xu W, Zhang Y, Lv F-J (2020) Clinical and immunological assessment of asymptomatic SARS-CoV-2 infections. Nature medicine 26: 1200–1204

Madhi SA, Baillie V, Cutland CL, Voysey M, Koen AL, Fairlie L, Padayachee SD, Dheda K, Barnabas SL, Bhorat QE et al (2021) Efficacy of the ChAdOx1 nCoV-19 Covid-19 Vaccine against the B.1.351 Variant. New England Journal of Medicine

McCray Jr PB, Pewe L, Wohlford-Lenane C, Hickey M, Manzel L, Shi L, Netland J, Jia HP, Halabi C, Sigmund CD (2007) Lethal infection of K18-hACE2 mice infected with severe acute respiratory syndrome coronavirus. Journal of virology 81: 813–821

Muyldermans S (2013) Nanobodies: natural single-domain antibodies. Annual review of biochemistry 82: 775–797

Nambulli S, Xiang Y, Tilston-Lunel NL, Rennick LJ, Sang Z, Klimstra WB, Reed DS, Crossland NA, Shi Y, Duprex WP (2021) Inhalable Nanobody (PiN-21) prevents and treats SARS-CoV-2 infections in Syrian hamsters at ultra-low doses. Science Advances 7: eabh0319

Planas D, Bruel T, Grzelak L, Guivel-Benhassine F, Staropoli I, Porrot F, Planchais C, Buchrieser J, Rajah MM, Bishop E (2021) Sensitivity of infectious SARS-CoV-2 B. 1.1. 7 and B. 1.351 variants to neutralizing antibodies. Nature medicine: 1–8

Ruetalo N, Businger R, Althaus K, Fink S, Ruoff F, Pogoda M, Iftner A, Ganzenmüller T, Hamprecht K, Flehmig B (2021) Antibody Response against SARS-CoV-2 and Seasonal Coronaviruses in Nonhospitalized COVID-19 Patients. Msphere 6: e01145–01120

Schepens B, van Schie L, Nerinckx W, Roose K, Van Breedam W, Fijalkowska D, Devos S, Weyts W, De Cae S, Vanmarcke S et al (2021) Drug development of an affinity enhanced, broadly neutralizing heavy chain-only antibody that restricts SARS-CoV-2 in rodents. bioRxiv: 2021.2003.2008.433449

Scudellari M (2020) How the pandemic might play out in 2021 and beyond. Nature: 22–25

Taylor A, Foo SS, Bruzzone R, Vu Dinh L, King NJ, Mahalingam S (2015) Fc receptors in antibody-dependent enhancement of viral infections. Immunological reviews 268: 340–364

Tegally H, Wilkinson E, Lessells RJ, Giandhari J, Pillay S, Msomi N, Mlisana K, Bhiman JN, von Gottberg A, Walaza S (2021) Sixteen novel lineages of SARS-CoV-2 in South Africa. Nature Medicine 27: 440–446

Tirado SMC, Yoon K-J (2003) Antibody-dependent enhancement of virus infection and disease. Viral immunology 16: 69–86

Traenkle B, Segan S, Fagbadebo FO, Kaiser PD, Rothbauer U (2020) A novel epitope tagging system to visualize and monitor antigens in live cells with chromobodies. Scientific reports 10: 1–13

Volz E, Mishra S, Chand M, Barrett JC, Johnson R, Geidelberg L, Hinsley WR, Laydon DJ, Dabrera G, O'Toole A et al (2021) Assessing transmissibility of SARS-CoV-2 lineage B.1.1.7 in England. Nature 593: 266–269

Wagner TR, Ostertag E, Kaiser PD, Gramlich M, Ruetalo N, Junker D, Haering J, Traenkle B, Becker M, Dulovic A et al (2021) NeutrobodyPlex-monitoring SARS-CoV-2 neutralizing immune responses using nanobodies. EMBO Rep 22: e52325

Wang P, Nair MS, Liu L, Iketani S, Luo Y, Guo Y, Wang M, Yu J, Zhang B, Kwong PD (2021) Antibody resistance of SARS-CoV-2 variants B. 1.351 and B. 1.1. 7. Nature 593: 130–135

Weinreich DM, Sivapalasingam S, Norton T, Ali S, Gao H, Bhore R, Musser BJ, Soo Y, Rofail D, Im J et al (2020) REGN-COV2, a Neutralizing Antibody Cocktail, in Outpatients with Covid-19. New England Journal of Medicine 384: 238–251

Winkler ES, Bailey AL, Kafai NM, Nair S, McCune BT, Yu J, Fox JM, Chen RE, Earnest JT, Keeler SP et al (2020) SARS-CoV-2 infection of human ACE2-transgenic mice causes severe lung inflammation and impaired function. Nat Immunol 21: 1327–1335

Wrapp D, De Vlieger D, Corbett KS, Torres GM, Wang N, Van Breedam W, Roose K, van Schie L, Covid V-C, Team R (2020) Structural basis for potent neutralization of betacoronaviruses by single-domain camelid antibodies. Cell 181: 1004–1015. e1015

Wu X, Cheng L, Fu M, Huang B, Zhu L, Xu S, Shi H, Zhang D, Yuan H, Nawaz W et al (2021) A potent bispecific nanobody protects hACE2 mice against SARS-CoV-2 infection via intranasal administration. bioRxiv: 2021.2002.2008.429275

Xiang Y, Nambulli S, Xiao Z, Liu H, Sang Z, Duprex WP, Schneidman-Duhovny D, Zhang C, Shi Y (2020) Versatile and multivalent nanobodies efficiently neutralize SARS-CoV-2. Science 370: 1479–1484

Xie X, Muruato A, Lokugamage KG, Narayanan K, Zhang X, Zou J, Liu J, Schindewolf C, Bopp NE, Aguilar PV (2020) An infectious cDNA clone of SARS-CoV-2. Cell host & microbe 27: 841–848. e843

Yong SJ (2021) Long COVID or post-COVID-19 syndrome: putative pathophysiology, risk factors, and treatments. Infectious Diseases: 1–18

Zhou D, Dejnirattisai W, Supasa P, Liu C, Mentzer AJ, Ginn HM, Zhao Y, Duyvesteyn HM, Tuekprakhon A, Nutalai R (2021) Evidence of escape of SARS-CoV-2 variant B. 1.351 from natural and vaccine-induced sera. Cell 184: 2348–2361. e2346

